# A systems biology analysis of lipolysis and fatty acid release from adipocytes *in vitro* and from adipose tissue *in vivo*

**DOI:** 10.1101/2020.12.18.423229

**Authors:** William Lövfors, Jona Ekström, Cecilia Jönsson, Peter Strålfors, Gunnar Cedersund, Elin Nyman

**Affiliations:** Department of Biomedical Engineering, Linköping University, Linköping, Sweden; Department of Mathematics, Linköping University, Linköping, Sweden; Department of Biomedical and Clinical Sciences, Linköping University, Linköping, Sweden; Center for Medical Image Science and Visualization (CMIV), Linköping University, Linköping, Sweden

**Keywords:** Systems biology, lipolysis, adipocytes, type 2 diabetes, signalling

## Abstract

Lipolysis and the release of fatty acids to supply energy fuel to other organs, such as between meals, during exercise, and starvation, are fundamental functions of the adipose tissue. The intracellular lipolytic pathway in adipocytes is activated by adrenaline and noradrenaline, and inhibited by insulin. Circulating fatty acids are elevated in type 2 diabetic individuals. The mechanisms behind this elevation are not fully known, and to increase the knowledge a link between the systemic circulation and intracellular lipolysis is key. However, data on lipolysis and knowledge from *in vitro* systems have not been linked to corresponding *in vivo* data and knowledge *in vivo*. Here, we use mathematical modelling to provide such a link. We examine mechanisms of insulin action by combining *in vivo* and *in vitro* data into an integrated mathematical model that can explain all data. Furthermore, the model can describe independent data not used for training the model. We show the usefulness of the model by simulating new and more challenging experimental setups *in silico*, e.g. the extracellular concentration of fatty acids during an insulin clamp, and the difference in such simulations between individuals with and without type 2 diabetes. Our work provides a new platform for model-based analysis of adipose tissue lipolysis, under both non-diabetic and type 2 diabetic conditions.

## Introduction

Lipolysis, the breakdown of triacylglycerol to glycerol and fatty acids, and the subsequent release of fatty acids and glycerol as energy fuel for other organs, is one of the main functions of the adipose tissue. Because of the critical role of fatty acids as a fuel for the body, this function is also central to energy homeostasis. Interest in lipolysis has gained more traction as the prevalence of obesity, type 2 diabetes and its sequelae have increased dramatically over the last decades. Lipolysis is stimulated in the body mainly by the catecholamine noradrenaline, which is released locally in the adipose tissue, and adrenaline, which is elevated in the circulation. The two catecholamines signal through α_2_- and β-adrenergic receptors. The beta-adrenergic receptors stimulate lipolysis by increasing intracellular levels of cyclic AMP (cAMP). An increased concentration of cAMP results in the activation of adipose triacylglycerol lipase (ATGL) and hormone sensitive lipase (HSL), the two rate-limiting lipases responsible for lipolysis. Insulin counteracts the stimulation of lipolysis in adipocytes by activation of phosphodiesterase 3B (PDE3B) that degrades cAMP and thereby reduces the rate of lipolysis [1–3]. The two catecholamines can also inhibit lipolysis by inhibiting the activation of adenylate cyclase through the α_2_-adrenergic receptor. Lipolysis is thus under tight positive and negative hormonal control.

The signalling pathways involved in the control of lipolysis are highly complex, and numerous crosstalks between different pathways and branches are emerging. Jönsson *et al*. [3] provide detailed elucidation of the pathways controlling lipolysis in adipocytes from human subcutaneous adipose tissue and show a new β-adrenergic – insulin crosstalk, where β-adrenergic signalling, in addition to stimulation, also inhibits lipolysis via parts of the insulin signalling pathway. They also demonstrate an additional stimulatory lipolytic action of insulin at high concentrations. Beyond the actions mentioned in [3], Stich *et al*. also suggest an anti-lipolytic action of insulin involving α_2_-adrenergic receptors [4]. This action was observed during microdialysis experiments, *in situ* in human subcutaneous adipose tissue, stimulated with adrenaline, isoproterenol, insulin and phentolamine. The high degree of crosstalk and the different actions at different concentrations of the hormones controlling lipolysis make it hard to successfully grapple with experimental data by mere reasoning. To understand the role and relative importance of these different actions of insulin and the catecholamines in the control of lipolysis, a next step therefore is to test the suggested mechanisms in a formalized way using mathematical modelling.

We have earlier, in several steps, developed mathematical models for insulin signalling in human adipocytes: first in isolation and later connected to models of systemic glucose control, and used the models to unravel key alterations in type 2 diabetes [5–8]. These models, however, do not include lipolysis and the control of lipolysis by insulin. We have also studied systemic whole-body effects of fatty acids on glucose uptake and release, using modelling [9], but with no link to intracellular lipolysis. There have also been other efforts to understand adipose tissue lipolysis in more detail, for example experimentally in [10] and using mathematical modelling in [11, 12], but without detailed intracellular components. In summary, none of the existing models have been developed to elucidate the mechanisms of control of intracellular lipolysis.

Here, we develop a new minimal model for lipolysis and the release of fatty acids based on both *in vitro* and *in vivo* experimental data from humans (Fig. 1). The model includes all three suggested insulin actions to control lipolysis: two direct actions, one anti-lipolytic via protein kinase B (PKB) activation of phosphodiesterase 3B (PDE3B) (action-1), one lipolytic via inhibition of PDE3B (action 2), and a third indirect anti-lipolytic action via α-adrenergic receptors (action-3). Using mechanistic modelling, we can evaluate the impact of these actions individually. The model accurately predicts independent validation data and is therefore useful to simulate new *in silico* experiments, such as the release of fatty acids *in vivo*, under both non-diabetic and type 2 diabetic conditions. The developed model is, to the best of our knowledge, the first model for the hormonal control of lipolysis, and it opens for new research and drug discovery related to type 2 diabetes.

**Figure 1:**
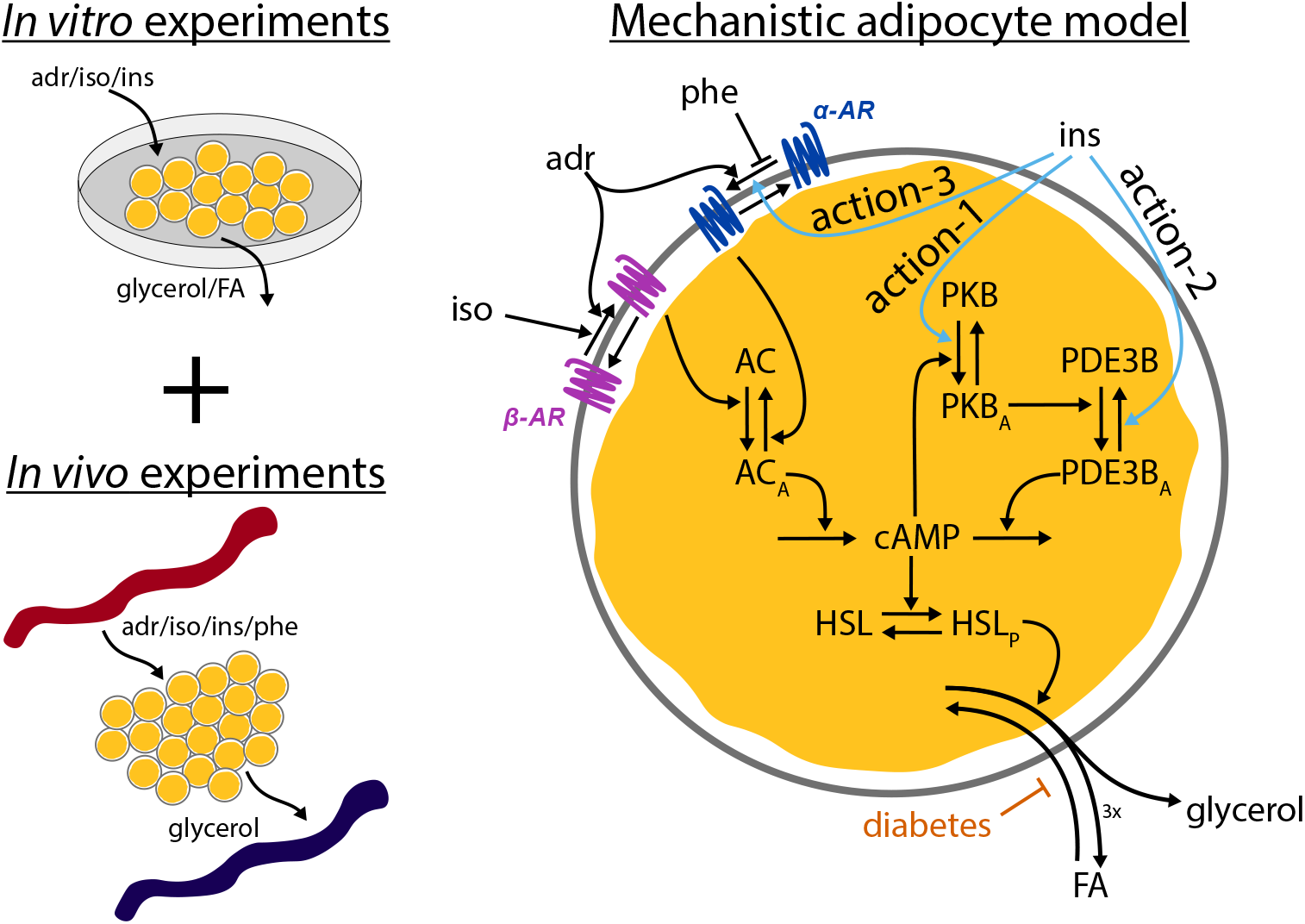
The system under study. Data from both *in vitro* and *in vivo* experiments of lipolysis control by insulin, adrenergic stimulus, and phentolamine were combined to create a first mechanistic model of lipolysis, under both non-diabetic and type 2 diabetic conditions. The model responds to stimuli with adrenaline (adr), isoproterenol (iso), insulin (ins), and phentolamine (phe), initiating signalling cascades through key proteins leading to release of fatty acids (FA) and glycerol. Adrenaline affects both β-adrenergic receptors (β-AR) and α-adrenergic receptors (α-AR), while iso only affects β-AR. Ins gives rise to three different insulin actions: action-1) an anti-lipolytic effect of insulin via protein kinase B (PKB) and phosphodiesterase 3B (PDE3B), action-2) a positive lipolytic effect via PDE3B at high insulin concentrations, and action-3) an anti-lipolytic effect of insulin via α-adrenergic receptors.

## Results

To connect data from several sources in a common framework, we use mechanistic modelling. In mechanistic modelling, available knowledge about a system is formulated as a model by constructing a set of ordinary differential equations. The validity of such models can be tested by comparing model simulations to experimental data. Typically, the values of the model parameters, e.g. kinetic rate constants and initial concentrations of substances, are unknown and need to be estimated by training the model to experimental data. Other experimental data are then used for testing and validating the predictive power of the model.

### Experimental observations and model development

We developed a mechanistic model focused on the regulation of intracellular lipolysis and the release of fatty acids and glycerol from the adipose tissue. The model is based on data from both *in vitro* measurements on isolated adipocytes, and *in vivo* microdialysis measurements, in both cases from non-diabetic individuals. More specifically, to develop the model we used experimental data from two sources: i) isolated adipocytes treated *in vitro* with the β-adrenergic agonist isoproterenol to stimulate lipolysis, and additionally with insulin to inhibit the isoproterenol-stimulated lipolysis, from [3], and ii) microdialysis measurements of lipolysis *in vivo*, stimulated with adrenaline/isoproterenol and inhibited by insulin and phentolamine, from [4]. We have taken three previously suggested mechanisms of crosstalk for the actions of insulin to explain the observed behaviour in the experimental data: 1) an anti-lipolytic effect of insulin via protein kinase B (PKB) and PDE3B [3, 13] – action-1, 2) a positive lipolytic effect of insulin via PDE3B at high concentrations of insulin [3] – action-2, and 3) an anti-lipolytic effect of insulin via α-adrenergic receptors [4] – action-3. We have also included other known signalling steps in the control of lipolysis in adipocytes as indicated in Fig. 1 and detailed in the Methods section. To avoid overfitting, the model was kept “minimal” in the sense that we focused on a few key proteins, and not every protein known to be involved in the control of lipolysis.

To further support the claim that the model is minimal, we performed a parameter identifiability analysis as detailed in the **Methods - Uncertainty estimation** section. Any rate-determining parameter (*k*_*x*_) that appeared to be (downwards) non-identifiable was removed from the model, with the exception of the parameter determining the reesterification (*k*8*c*) since that parameter was necessary to implement the effect of the diabetic condition later. The parameter uncertainty bounds for the final minimal model is shown in Fig. 2 and Table S1. In total, 23 parameters were found to be identifiable.

**Figure 2:**
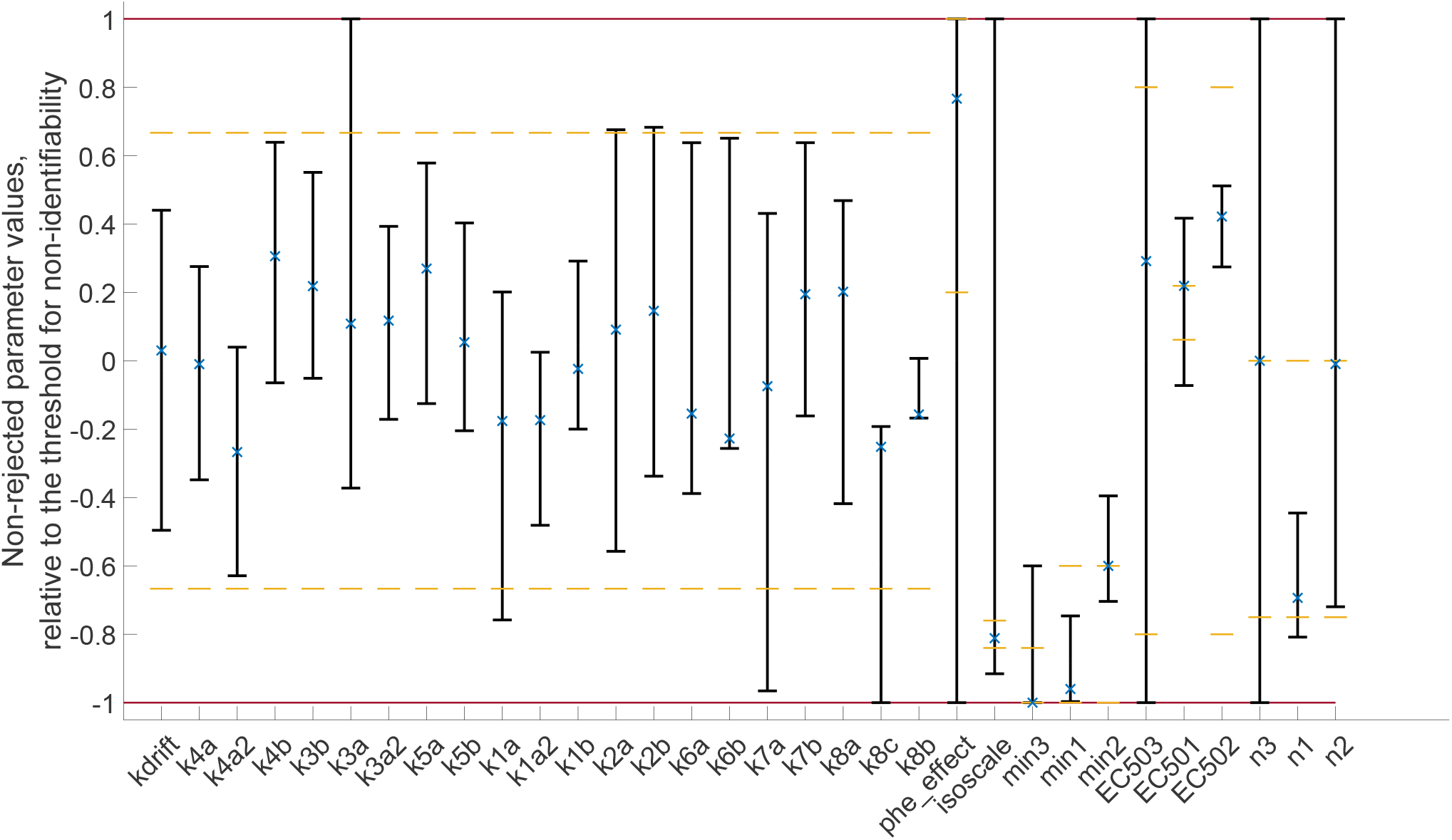
Parameter identifiability analysis. The minimal and maximal values of a parameter was found using the optimization approach detailed in the **Methods - Uncertainty estimation** section. The parameter values are expressed as relative values with respect to threshold for non-identifiability, where -1 represents a parameter value at the lower threshold, and 1 represents a parameter at the upper threshold. x represent the optimal parameter values and yellow dashed lines represent the bounds for a specific parameter when estimating parameter values. Bounds, thresholds and parameter values are shown in Table S1 and Table S2.

### Comparisons between model simulations and data

The model was trained to *in vitro* dose-response data for the phosphorylation of PKB at Ser-473 and the release of glycerol and fatty acids in response to isoproterenol and insulin stimulation, as well as *in vivo* microdialysis data of glycerol release in response to adrenaline and insulin (Fig. 3; solid lines represent the model simulation with the best agreement to data, shaded areas represent the model uncertainty, and experimental data are represented as mean values with error bars (SEM)). Here, the model uncertainty refers to the most extreme simulations, while still requiring the model to pass a *χ*^2^-test. The best model simulation clearly has a good agreement with the experimental data (Fig. 3). This visual assessment is supported by a statistical *χ*^2^-test, where the cost of the model (*v*^*^ for the optimal cost, see **Methods**), given the optimal parameter values (*θ*^*^, see Table S2), is below the threshold of rejection given by the *χ*^2^-test (*v*^*^ = 130.8 < *χ*^2^(0.05, 137) = 165.3) for a significance level of 0.05. The optimal parameter values are shown in Table S2 and in the set of scripts used to reproduce the results (see **Data and model availability**). The model uncertainty was estimated in the same way as in [14], by maximizing/minimizing the simulation in all experimental data points while requiring the cost to not exceed the *χ*^2^-threshold for a significance level of 0.05.

**Figure 3:**
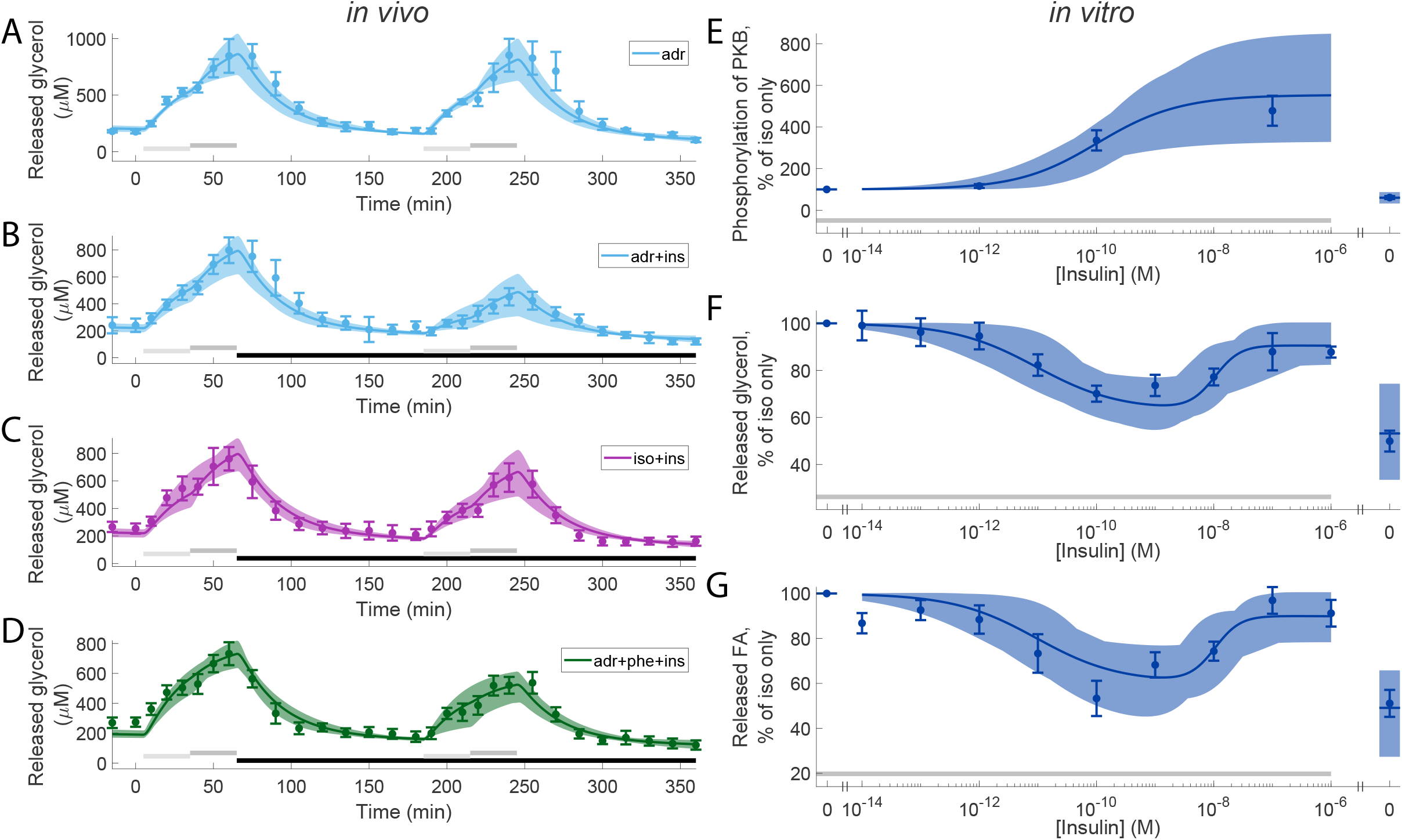
Model agreement with experimental data. In all panels, solid lines represent the model simulation with the best agreement to data, the shaded areas represent the model uncertainty, and experimental data points are represented as mean values with error bars (SEM). (A-D), *in vivo* time-series experiments. (E-G),*in vitro* dose-response experiments. In all subfigures, horizontal bars indicate where stimulations were given. In detail, light/dark grey bars indicate stimulation with: 1/10 µM, respectively, adrenaline in (A,B), 0.1/1 µM isoproterenol in (C), and 1/10 µM adrenaline with 100 µM phentolamine. Black bars in (B-D) indicates stimulation with 0.6 nM insulin. In (E-G) grey bars indicate stimulation with isoproterenol (10 nM). In the *in vivo* experiments, experiments with adrenaline are shown in light blue (A-C), with isoproterenol in purple (B), and with the combined stimulation with adrenaline and phentolamine in green (C). In the *in vitro* experiments (D-F), increasing doses of insulin were given together with 10 nM isoproterenol in all points except one. The point without isoproterenol got no stimulus and is shown to the right in the graphs. An alternative visualization is available in Fig. S1 show the difference by overlaying the experiments.

### Model testing: predicting intracellular phosphorylation of HSL

For a model to be of practical use, it should be able to perform reasonable predictions. To test this, we used the model to predict the dose-response for phosphorylation of HSL (HSLp), an intracellular state in the model (Fig. 1) that was not used when training the model to data. We estimated the uncertainty of the model prediction in the same way as described in the comparison between model and data, i.e. we maximized/minimized the prediction simulation, while requiring the agreement to the estimation data to be acceptable (keeping the cost below the threshold, see **Statistical analysis** in **Methods**). When compared to the experimental data (Fig. 4), the model prediction overlaps well. This is statistically supported using a *χ*^2^-test (*v*^*^ = 10.7 < *χ*^2^(0.05, 10) = 18.3).

**Figure 4:**
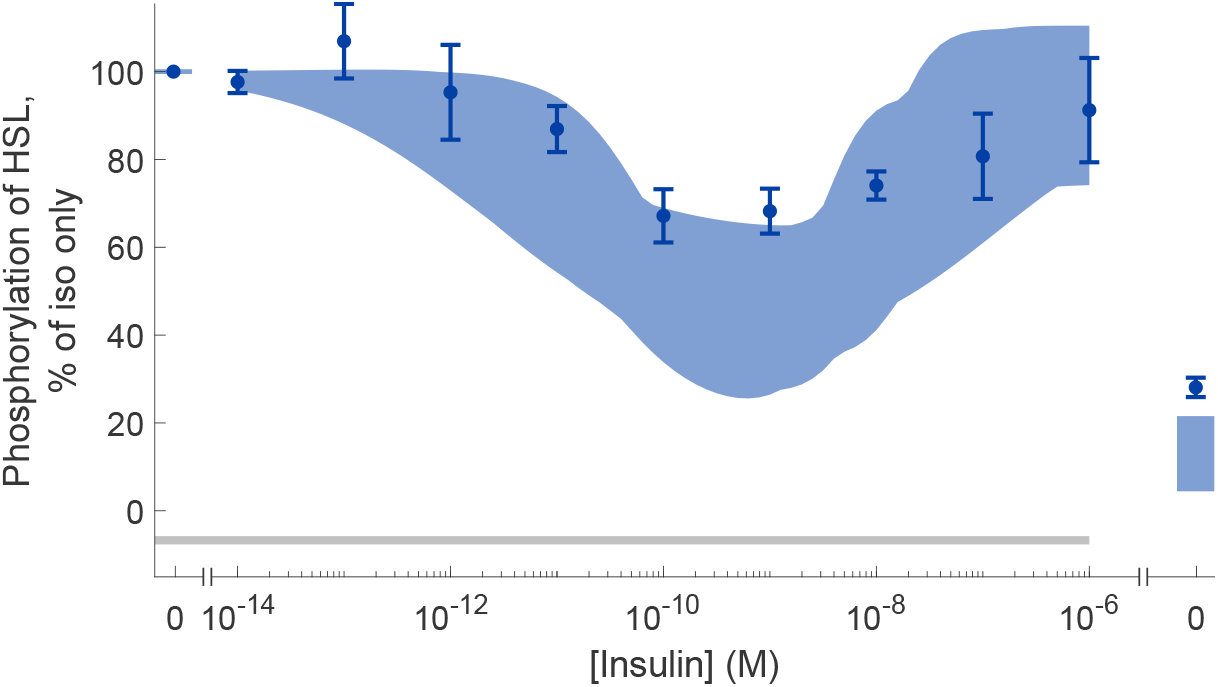
Prediction of intracellular extent of HSL phosphorylation. The shaded area represents the model uncertainty, and the experimental data are represented as mean values with error bars (SEM). The horizontal grey bar indicates where stimulation with isoproterenol (10 nM) have been given. Increasing doses of insulin were given together with 10 nM isoproterenol in all points except one. The point without isoproterenol got no stimulus and is shown to the right in the graph.

### Investigating the different actions of insulin

With the validated model, we continued to investigate the impact of the three different insulin actions (Fig. 1, the three blue arrows) by excluding one action at a time. We excluded an action by keeping the corresponding *Ins*_*x*_ variable (see Eq. (1)) at basal levels throughout the simulation (instead of increasing with increased concentration of insulin). Firstly, by removing action-1 (the antilipolytic effect of insulin via PKB-mediated activation of PDE3B), the model is unable to explain the decline in glycerol release in response to increased levels of insulin *in vitro*: compare Fig. 5A with Fig. 5C. Secondly, by removing action-2 (the positive lipolytic effect of insulin via inhibition of PDE3B), the model is unable to explain the recovery in glycerol release at high insulin concentrations *in vitro*: compare Fig. 5A with Fig. 5E. Finally, by removing action-3 (the anti-lipolytic effect of insulin via α-adrenergic receptors), the model is unable to explain the decrease in glycerol release in the second set of adrenergic stimuli *in vivo*: compare Fig. 5B with Fig. 5D. The removal of insulin action-1 and -2 renders the model unable to agree with the experimental data sufficiently well 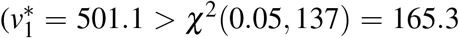 and 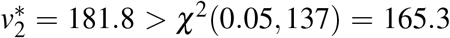 for the removal of action-1 and -2 respectively). With the removal of action-3 the model can still quantitatively explain the data sufficiently well 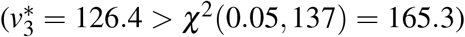, but not qualitatively. In the experimental data, the release of glycerol is similar when comparing the first stimulation with adrenaline to the stimulation with both adrenaline and phentolamine. In the second stimulation with both adrenaline and insulin (and phentolamine), the release of glycerol is markedly lower when phentolamine is present. This decrease in the release of glycerol is absent in simulations of the model without insulin action-3.The best agreement between the model without insulin action-3 and experimental data is shown in Fig. S2. Consequently, all three actions of insulin are required for the model to explain the available experimental data.

**Figure 5:**
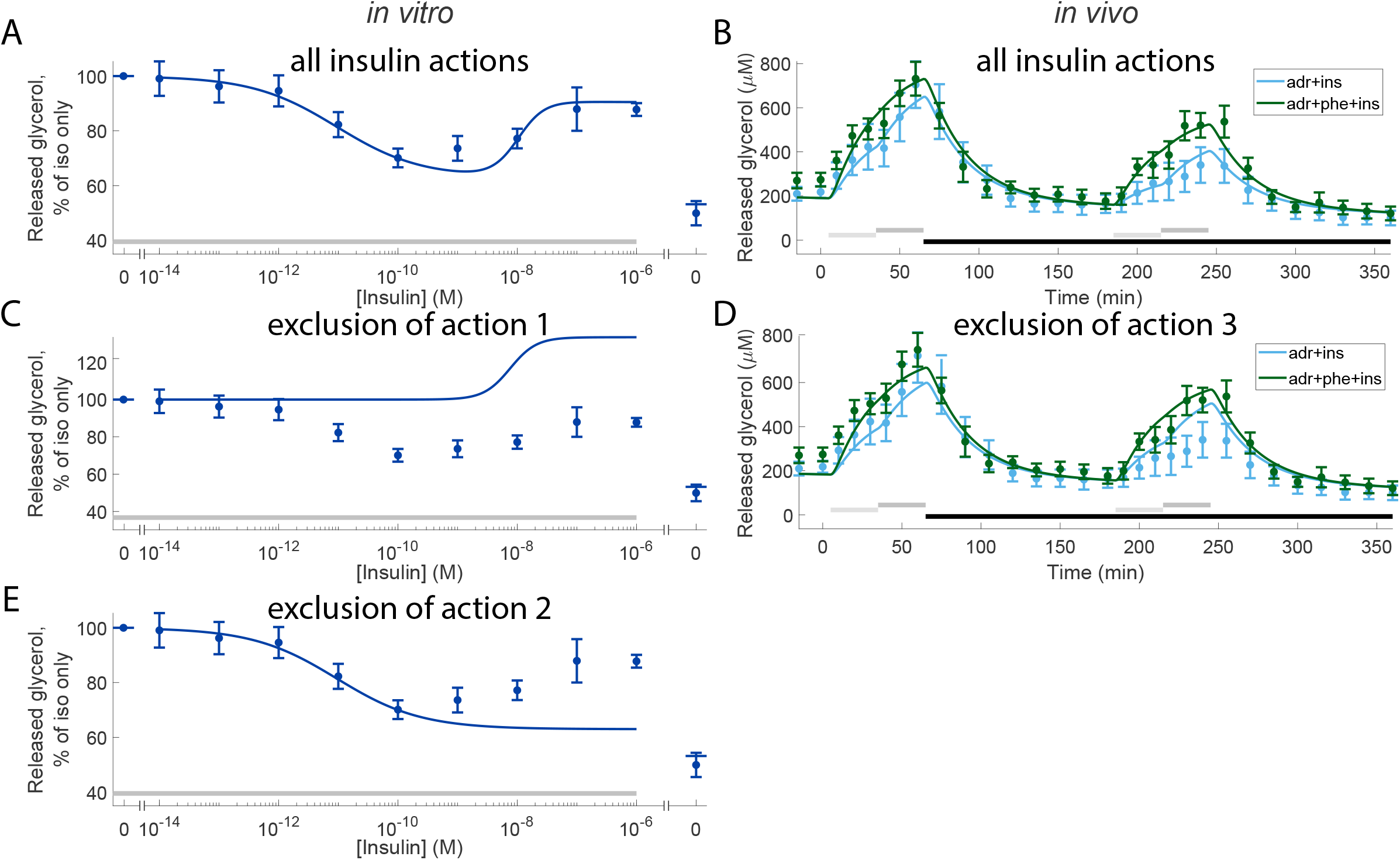
Effects of excluding either of the three insulin actions. In panels (A-E), data points with error bars represent mean and SEM values and solid lines represent the model simulation with the best agreement to data. (A, B), model simulations with all insulin actions present (same as Fig. 3C, F), see Fig. 1 for a graphical representation of the three insulin actions. (C-E), the model simulations when either of the actions are excluded. In all subfigures, horizontal bars indicate where stimulations were given. In (A, C, E) grey bars indicate stimulation with isoproterenol (10 nM) and in (B, D) light/dark grey bars indicate stimulation with low/high dose of adrenaline (1/10 µM) with or without phentolamine (100 µM), and black bars indicate stimulation with insulin (0.6 nM). In the *in vitro* experiments (A, C, E), increasing doses of insulin were given together with 10nM isoproterenol in all points except one. The point without isoproterenol got no stimulus and is shown to the right in the graphs. In the *in vivo* experiments, experiments with adrenaline are shown in light blue (B, D) and with the combined stimulation with adrenaline and phentolamine in green (D).

### Estimating the extent of altered reesterification in type 2 diabetes

We then used the validated model to gain new biological insights. As demonstrated in [3, Fig. 14] at the cellular level, essentially only the release of fatty acids, but not of glycerol and hence not lipolysis, is affected in type 2 diabetes. The authors conclude that this is due to reduced reesterification, i.e. a decreased reuse of fatty acids to re-form triacylglycerol, in the diabetic state. We therefore added a single parameter representing a decrease in reesterification to the model to represent the type 2 diabetic condition (*diab* in Eq. (9)). In addition to extending the model, we also extended the set of experimental data beyond the data used so far (see e.g Fig. 3). The set of experimental data now also includes the phosphorylation of HSL previously used for validation (Fig. 4), as well as reesterification under type 2 diabetic conditions ([3, Fig. 14F]). We then trained the extended model with the extended set of experimental data and quantified the maximal range of reesterification under both normal and type 2 diabetic conditions. The model agrees well with the experimental data (Figs. 6 and S3), supported by a *χ*^2^-test (*v*^*^ = 164.1 < *χ*^2^(0.05, 152) = 181.8), and accurately shows that only the release of fatty acids and thus reesterification is affected under type 2 diabetic conditions (Fig. 6). With the trained model, we found the range of reesterification to be altered from 66.7 - 74.3 % under normal conditions to 39.6 - 64.1% under diabetic conditions.

**Figure 6:**
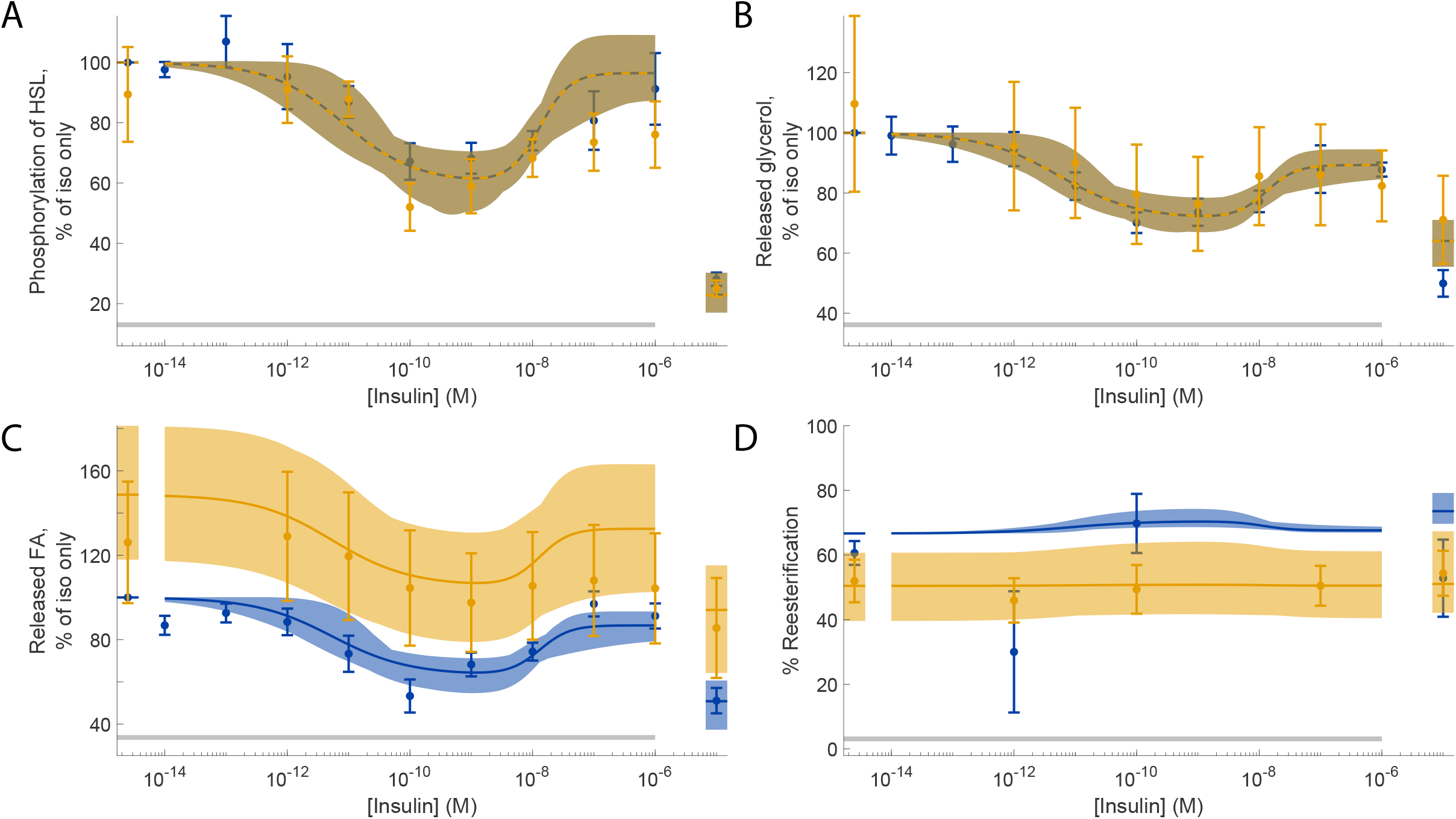
Model agreement when trained to the extended set of experimental data. In all panels, dots and error bars represent mean and SEM values, solid/dashed lines represent the model simulation with the best agreement to data, and the shaded areas represent the model uncertainty. (A), phosphorylation of HSL. (B), release of glycerol. (C), release of fatty acids (FA). (D), percentage of fatty acids being reesterified. Blue data/simulations correspond to normal conditions, and orange data/simulations correspond to type 2 diabetic conditions. The data for diabetic conditions in (A-C) and the data for normal conditions (D) were not used to train the model. In all panels, horizontal grey bars indicate where stimulation with isoproterenol (10 nM) was given. Furthermore, increasing concentrations of insulin were given together with 10nM isoproterenol in all points except one in all panels. The point without isoproterenol got no stimulus and is shown to the right in the graphs. The agreement with the rest of the original dataset (*in vitro* experiments) is shown in Fig. S3.

### *In vivo* model simulations of fatty acid release, under both non-diabetic and type 2 diabetic conditions

In addition to predicting dose-response data or quantifying the range of impairment of the reesterification of fatty acids, we can also use the model to predict temporal changes *in vivo*. During lipolysis both glycerol and fatty acids are released from the adipocytes (Fig. 1). However, only the time series for glycerol release were measured in the *in vivo* data used to train the model parameters. We can now use the model to not only predict the release of fatty acids *in vivo* in response to e.g. treatment with adrenaline, we can also predict the fatty acid release *in vivo* under diabetic conditions. As expected, the release of fatty acids *in vivo* in response to adrenaline temporally mimics the release of glycerol (Fig. 7, cf. Fig. 3). Furthermore, in line with the finding that the reesterification is impaired under the diabetic condition, resulting in an increased release of fatty acids from the adipocytes, the model predicts that the adipose tissue release of fatty acids *in vivo* is increased under diabetic conditions.

**Figure 7:**
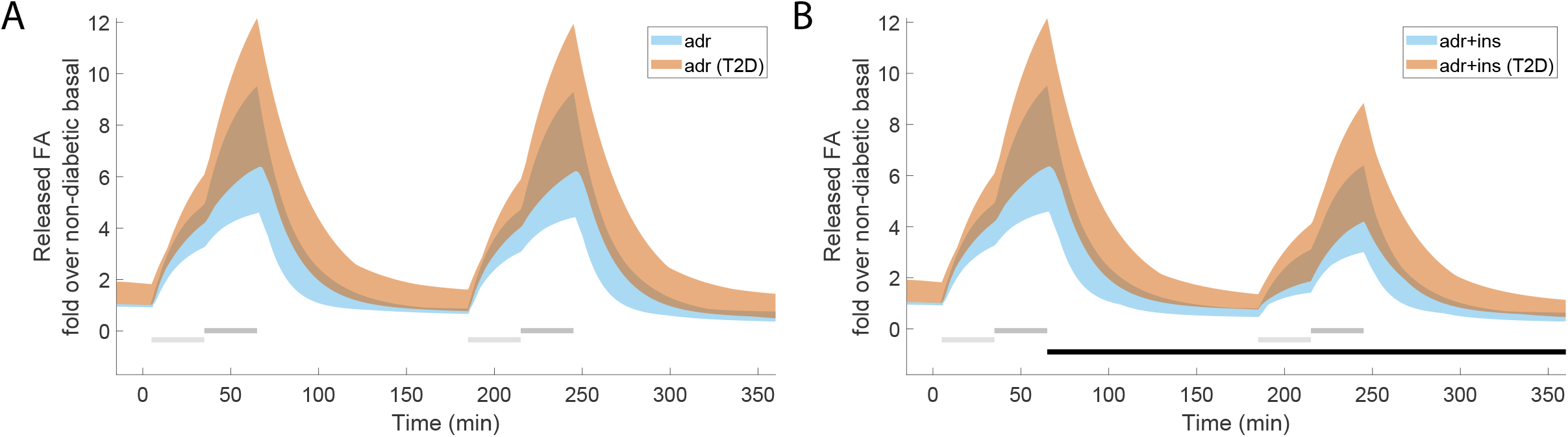
Simulations of fatty acid release *in vivo*, under both non-diabetic (blue) and type 2 diabetic conditions (orange). In both panels, the shaded areas represent the model simulation uncertainty. In both panels (A-B) horizontal light grey bars indicate where stimulation with 1 µM adrenaline was given, dark grey bars indicate where stimulation with 10 µM adrenaline was given, and horizontal black bars indicate where stimulation with 0.6 nM insulin was given. (A), model prediction without insulin, and (B), with insulin.

## Discussion

We examined the role of adipose tissue in the storage and release of fuel in the form of fat under normal conditions and when disturbed by insulin resistance and type 2 diabetes. We present a mathematical model of hormonal control of lipolysis in human adipose tissue (Fig. 1), a model that links molecular events at the cellular level with corresponding responses at the tissue level. The model can explain both *in vivo* temporal data from microdialysis experiments in adipose tissue (Fig. 3A-C and Fig. S3A-C) and *in vitro* dose-response data from isolated adipocytes (Fig. 3D-F, Fig. 6, and Fig. S3D-F), as well as accurately predict independent validation data (Fig. 4).

There exist other models of lipolysis in humans. For example [15] modelled insulin levels and fatty acid release in response to glucose intake on a systemic level, but do not model any adrenergic stimulus or have a detailed intracellular compartment. A more extensive model of lipid metabolism [12], is also missing a detailed intracellular compartment and adrenergic stimulus. Conversely, [16] modelled lipolysis in response to adrenergic stimulus with a detailed intracellular compartment, but lacked insulin signalling. Tangentially, there also exist models of glucose homeostasis with insulin signalling, but lacking both lipolysis and adrenergic hormonal control [7, 17]. The model of hormonal control of lipolysis presented here is the first, to our knowledge, that includes insulin and adrenergic signalling, as well as an intracellular compartment. Some of the existing models are more detailed for certain aspects of lipolysis. We have chosen to only include sub-systems directly supported by experimental data, and the presented model can therefore be considered ”minimal”.

The presented model is in agreement with the experimental data (Figs. 3, 6 and S3). Furthermore, the estimated uncertainty of the model is reasonably large compared to data uncertainty. This indicates that we have been successful in estimating the uncertainties of the model parameters and simulations.

Insulin action-3 – the anti-lipolytic effect of insulin via α-adrenergic receptors – was needed to explain the combined effect of adrenaline, insulin, and phentolamine seen in the *in vivo* estimation data, as become clear when the action was removed (Fig. 5D, notably in the second peak). It should be noted that insulin action-3, via the α-receptor, is also only observed at high concentrations of insulin, and may therefore be a secondary effect elicited by insulin in other cells or tissues.

In addition to investigating the contributions of the different insulin actions, we used the model to examine changes under type 2 diabetic conditions. The model shows that the reesterification is altered from 66.7 - 74.3 % under normal conditions to 39.6 - 64.1 % under type 2 diabetic conditions. We also used the model to predict the temporal release of fatty acids *in vivo* in response to adrenaline in both non-diabetic and type 2 diabetic conditions (Fig. 7). Type 2 diabetes and obesity have traditionally been associated with elevated levels of circulating fatty acids, an issue both challenged and affirmed [10, 18]. Our model predicts an *in vivo* increase in the release of fatty acids in diabetic conditions versus in non-diabetic conditions. However, our model does not include mechanisms for fatty acid clearance due to uptake by other organs and therefore cannot predict the systemic levels of fatty acids or possible changes to the clearance in type 2 diabetes.

We used data from two different sources [3, 4] to develop the model, which can be seen both as a strength and as a weakness. The use of internally consistent data, from the same laboratory under the same experimental conditions, is potentially important to test hypotheses to unravel new biological mechanisms. For the purpose herein, to develop a first intracellular model of lipolysis that includes key observations and that later can be further built upon when more data become available, we believe it is a strength to use data from multiple sources. This means that the model is more general, and therefore more likely to be useful together with other human data from studies of adipose tissue lipolysis.

Desensitization, that cells decrease their response to continued or repeated stimuli with time, is a known phenomenon of β-adrenergic signalling. Stich et al [4] controlled for desensitization by using multiple repeats of the stimulus paradigm. We decided to only include the first two rounds of stimulation of lipolysis, as we think there is a tendency to desensitization in the third stimulation, and we decided not to include this behaviour in this first model. There are other studies that show a clear desensitization in the release of glycerol in response to adrenaline [19, 20]. These studies have shorter intervals between stimuli (30 min, 1 h), and also reveal desensitization due to exercise-induced stimulation of lipolysis [20]. We have previously studied desensitization in heart cells using modelling and found that dose-response curves need to be adjusted for this phenomenon before important parameters such as the EC50 are computed [21]. Desensitization is an important phenomenon in β-adrenergic signalling that should not be overlooked and should be addressed in later models of lipolytic control.

Furthermore, in the *in vivo* data there were two different sets of data from the same experimental condition (adrenaline with insulin, Figs. 2A and 3A in [4]). Since the model can only produce one simulation for identical conditions, we decided to not use two datasets for identical conditions to compute the cost. At the same time, we needed to scale the model states to be comparable to the experimental data. Therefore, we used the adrenaline with insulin data from Fig. 2A in [4] to both compute optimal scaling parameters and the cost of the model, and only used the data from Fig. 3A in [4] (shown in Fig. S1C) to compute scaling parameters.

During the modelling, we constrained insulin action-2 (Fig. 1) so that the EC50 in response to insulin stayed between 0.5 and 1.1 nM. The reasoning behind this constraint is that known upstream signalling intermediaries, such as the autophosphorylation of the insulin receptor, has an EC50 in that range [5]. We also included a slightly delayed response to changes in adrenaline and isoproterenol stimulus *in vivo* when developing the model. Such a delay was observed in the microdialysis data [4], but not as obviously present in primary adipocytes [3]. In effect, we delayed the time for all changes of adrenaline and isoproterenol concentrations with 5 min when simulating the *in vivo* data (e.g. in Fig. 3A-C). The agreement between model simulations and data became substantially better with this delay. The underlying cause behind the observed delay in the microdialysis setup could be pooling of data or a delay in the microdialysis probe – or a biological tissue effect. We have chosen not to include the mechanisms of the delay in the model, instead we explicitly added the time delay.

The model developed herein can be further developed in several directions. First, on the intracellular side, we aim to combine the model with our extensive work on the modelling of insulin signalling pathways in both non-diabetic and type 2 diabetic conditions [5–8]. With such a connection, we will reach a first comprehensive model for the human adipocyte that is based on extensive data from both non-diabetic and type 2 diabetic patients. Such a model would be able to simulate the major functions of the adipocyte: the control of lipolysis, as well as insulin control of glucose uptake, protein synthesis, and transcription. Second, on the systemic regulation of fatty acid release, the work herein opens up for a first connected model where intracellular components of lipolysis are connected to whole-body changes in fatty acid release. This connected model can also be combined with other models, for example models for intake of meals. Such a connected model is key to understand mechanisms of ectopic fat storage, i.e. where liver and muscle tissue increase their storage of lipids – a condition that is linked to disease development such as in type 2 diabetes, liver failure, and cardiovascular disease [22].

## Methods

### Data processing

*In vitro* experiments with isolated human adipocytes were performed by us and were previously published (Fig. 14D in [3]). In the *in vitro* data, two points (for 10^−7^ and 10^−6^ M insulin in the non-diabetic fatty acid release data) had only two repeats, and we therefore set the data uncertainty for those two points to the average uncertainty of the non-diabetic fatty acid release. *In vivo* data from human microdialysis experiments were extracted from Figs. 1A, 2A, and 3A in [4].

In all *in vivo* experiments [4], three sets of consecutive adrenergic stimuli were given. We have chosen to only include the first two sets of stimuli in the present study. The third set of stimuli was essentially a repeat of the second set of stimuli at a later time point - yet showed a different response than the second set of stimuli (not shown). The difference in the response can be technical and/or biological differences at this later time, differences not included in the model in the present study. Furthermore, we used only one of the two datasets with identical stimuli (adrenaline and insulin) for the calculation of the cost. However, we kept the other dataset when calculating the experimental scaling parameters. Details of these decisions are discussed in the Discussion section.

### Mathematical modelling

A system of ordinary differential equations (ODEs) was used to model lipolysis and the release of fatty acids *in vitro* and *in vivo*. The equations are visualized in the interaction graph in Fig. 1. The full model with all described equations can be found in the supplementary files (see **Data and model availability**).

### Equations for insulin actions

We modelled the three different actions of insulin described in [3, 4] (see Fig. 1) as three separate sigmoidal functions, all dependent on the concentration of insulin. These three insulin functions affect downstream signalling proteins on PKB, PDE3B and the α_2_-adrenergic receptor. The equation for these sigmoidal functions is described in Eq. (1).

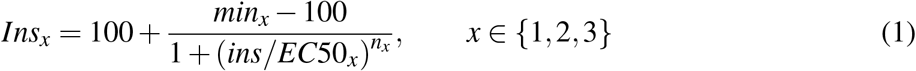

*Ins*_*x*_ is a function that is dependent on the given concentration of insulin (*ins*); *min*_*x*_ is the minimum value the function has at ins=0; *EC*50_*x*_ is the concentration of insulin at which the function reaches half of the maximal response (which is set to 100). The steepness of the function is determined by *n*_*x*_.

### Equations for insulin signalling

When IR is stimulated with insulin, a signalling cascade is initiated which leads up to the activation of PKB, through insulin receptor substrate 1 (IRS1), phosphoinositide 3-kinase (PI3K), and Phosphoinositide-dependent kinase-1 (PDK1). In our model, we have simplified this cascade as a direct action from insulin to the activation of PKB. PKB can also be activated by cAMP. We model PKB as either being in an inactive or an active configuration. The ODEs for PKB are given by Eq. (2)

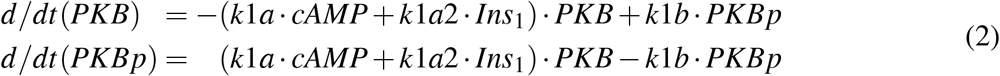

Here, *PKB* and *PKBp* are the two model states for the inactive and active form of PKB, *Ins*_1_ is the effect of insulin action-1, *cAMP* is the levels of cAMP, and *k*1*a k*1*a*2, and *k*1*b* are rate determining parameters.

Downstream, PKB will directly activate PDE3B. Beyond the direct activation by PKB, PDE3B will also be inactivated by insulin action-2. The detailed mechanism for this activation is currently unknown. We model PDE3B as being in either an inactive or an active state. The ODEs for PDE3B are given in Eq. (3).

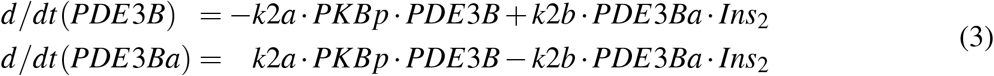

Here, *PDE*3*B* and *PDE*3*Ba* are the two model states for the inactive and active form of PDE3B, *PKBp* is the model state for activated PKB, *Ins*_2_ is the effect of insulin action-2, and *k*2*a* and *k*2*b* are rate determining parameters.

### Equations for α_2_-adrenergic receptor signalling

In the model, the α_2_-adrenergic receptor can switch between two different configurations: activated or inactivated. This balance is offset towards the activated state when the α_2_-adrenergic receptor is stimulated with adrenaline. The activated receptor will passively return to the inactive configuration. The activation of the α_2_-adrenergic receptor is also augmented by insulin and inhibited by the addition of phentolamine. The ODEs for the α_2_-adrenergic receptor are given in Eq. (4).

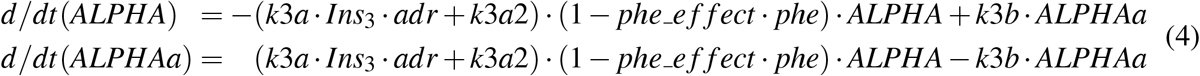

Here, *ALPHA* and *ALPHAa* are the two model states for the inactive and active α_2_-adrenergic receptor, *adr* and *phe* are the stimulation given as inputs, *Ins*_3_ is insulin action-3, *k*3*a, k*3*a*2 and *k*3*b* are rate determining parameters, and *phe e f f ect* is a parameter determining the effect of the phentolamine stimulation. In practice, *adr* corresponds to the concentration of adrenaline (in nM), and *phe* is a simplified boolean input (set to 1 if phentolamine is present, and 0 else).

### Equations for β-adrenergic receptor signalling

The β-adrenergic receptor, similarly to the α_2_-adrenergic receptor, is also switching between two configurations. In contrast to the α_2_-adrenergic receptor, the β-adrenergic receptor is also activated by adrenaline. Due to the uncertainty in difference in activation between adrenaline and isoproterenol, we added a scaling factor on isoproterenol. Furthermore, the activation of the β-adrenergic receptor is not increased by insulin or inhibited by phentolamine. The ODEs for the β-adrenergic receptor are given in Eq. (5).

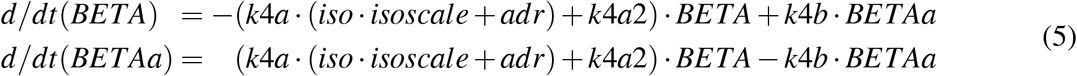

Here, *BETA* and *BETAa* are the two model states for the inactive and active β-adrenergic receptor, *adr* and *iso* are inputs corresponding to stimulation with adrenaline and isoproterenol, and *k*4*a, isoscale, k*4*a*2, and *k*4*b* are rate determining parameters. *adr* and *iso* corresponds to the concentration (in nM) of adrenaline and isoproterenol respectively.

### Equations for lipolysis

Both the β-adrenergic receptor and the α_2_-adrenergic receptor are G-protein coupled receptors. The G-proteins is made of multiple subunits, which will disassociate when the receptors are activated. These subunits will then go on and trigger another downstream effector. One such effector is adenylyl cyclase, which catalyse the conversion of ATP into cAMP. Through the G-proteins, β-adrenergic receptor will increase the activity of adenylyl cyclase, and α_2_-adrenergic receptor will decrease the activity. In the model, we have simplified this interaction by ignoring the G-proteins, letting the β-adrenergic receptor and α_2_-adrenergic receptor directly affect the adenylyl cyclase. Furthermore, we model the adenylyl cyclase as being either inactive or active, where the active version leads to an increased production of cAMP. The model equations are given in Eq. (6).

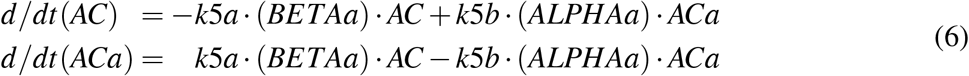

Here, *AC* and *ACa* are the two model states for inactive and active adenylyl cyclase respectively, *BETAa* and *ALPHAa* are the model states for active β- and α-receptor, and *k*5*a* and *k*5*b* are rate determining parameters.

Downstream of both adenylyl cyclase and PDE3B is cAMP. An increase in adenylyl cyclase activation will lead to an increased concentration of cAMP, and an increase in PDE3B activation will lead to a decreased concentration of cAMP. Together, adenylyl cyclase and PDE3B balance the concentration of cAMP in the cell. cAMP have negative feedback loop by activating PKB via PI3K, which in turn activates PDE3B, which leads to a decreased concentration of cAMP. We have also added both a basal production and degradation of cAMP. The ODEs for cAMP are given in Eq. (7).

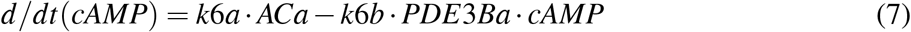

Here, *cAMP* is the model state for cAMP, *ACa* and *PDE*3*Ba* are the model states for the active configurations of adenylyl cyclase and PDE3B, and *k*6*a* and *k*6*b* are rate determining parameters.

The concentration of cAMP indirectly controls the lipolysis by activation of protein kinase A (PKA), which in turn will activate the lipid droplet-coating protein perilipin 1 (PLIN1) and hormone-sensitive lipase (HSL). Activation of PKA leads to the phosphorylation of the lipid droplet-coating protein perilipin 1 (PLIN1) and its subsequent release of the adipose triacylglycerol lipase (ATGL) activator, comparative gene identification-58. Active ATGL will catalyse the hydrolysis of the first fatty acid of the triacylglycerol. Phosphorylation of HSL by PKA results in activation and translocation of HSL to the lipid droplet, where HSL hydrolyses the second fatty acid leaving monoacylglycerol to be hydrolysed by a constitutively active monoacylglycerol lipase. HSL is capable to hydrolyse triacylglycerol, but ATGL is believed to be more important in this rate-limiting step of lipolysis. In the model, we simplified these interactions by only modelling HSL with an input from cAMP. The ODEs are given in Eq. (8).

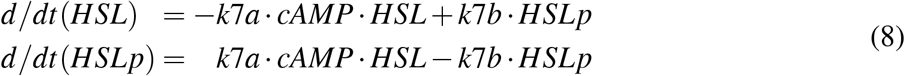

Here, *HSL* and *HSLp* are the states for inactive and active HSL, *cAMP* is the model state for cAMP, and *k*7*a* and *k*7*b* are the rate determining parameters.

Once the triacylglycerol has been broken down into three fatty acids and one glycerol, the glycerol will be transported out of the cell, and the fatty acids will either be transported out or reesterified with glycerol-3P into new triacylglycerol. This reesterification is reduced in type 2 diabetes. Some of the fatty acids can also go back into the cell, while the glycerol cannot. Fatty acids and glycerol outside of the cell will be cleared in an *in vivo* setting, but not *in vitro*. The ODEs are given in Eq. (9).

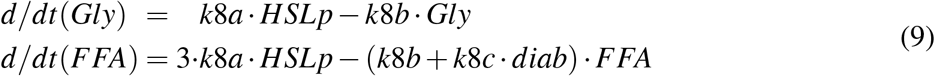

Here, *Gly* and *FFA* are the states for glycerol and fatty acids, *HSLp* is the state for activated HSL, *diab* is a parameter controlling the reduction of reesterification under diabetic conditions, and *k*8*a, k*8*b*, and *k*8*c* are rate determining parameters. *k*8*a* corresponds to the transport of fatty acids and glycerol from the inside to the outside of cell, *k*8*b* is the clearance of fatty acids and glycerol *in vivo* (clearance was disabled *in vitro* by setting *k*8*b* = 0), and *k*8*c* is the reesterification of fatty acids into triacylglycerol. To simulate the effect of type 2 diabetes on the reesterification, the parameter *diab* was allowed to vary between 0.0 – 1.0. In non-diabetic conditions, the type 2 diabetes effect was disabled by setting *diab* = 1 (i.e. no effect of type 2 diabetes).

### Translating the model states to experimental data

We constructed measurement equation to translate the model states of our model to the corresponding *in vivo* experimental data. In practice, we introduced a linear drift, a scaling constant and an offset constant. The measurement equation for glycerol is illustrated in Eq. (10).

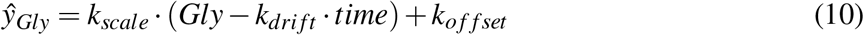

Here, *Gly* and *ŷ*_*Gly*_ are the model state and measurement equation for glycerol, *k*_*drift*_ ·*time* is the drift over time, *k*_*scale*_ is the scaling constant and *k*_*o ffset*_ is the offset constant. The scaling and offset constants were calculated using MATLABs least squares with known covariance (lscov) function.

For the *in vitro* experiments we did not use a measurement equation, but we did scale the simulations to be “fold over iso stimulation only”, as was done in the experimental data.

### Initial values

All states corresponding to activations were represented as per cent of activation, i.e. the sum of the two states will be 100. All initial values of the ODEs were set to arbitrary non-negative values:

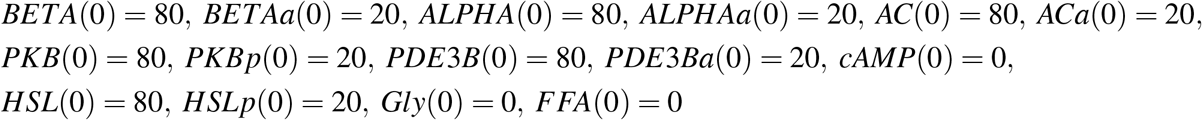

We then simulated the model without any stimuli to numerically calculate the steady state, which was used as initial values for the simulations of the experiments with stimuli.

### Calculating the percentage of reesterification

We calculate the percentage of reesterification using Eq. (11), the same way as the calculation was done for the experimental data in [3].

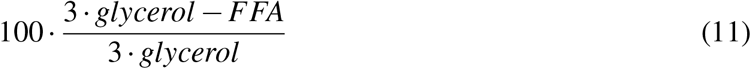

### Quantifying the model agreement to experimental data

In order to evaluate the model’s performance, we quantified the model agreement to data using a function typically referred to as a cost function. In this case, we used the normalized sum of squared residual as cost function (Eq. (12)).

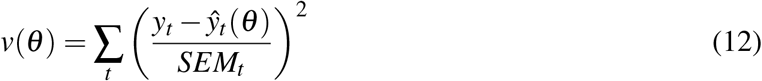

Here, *v*(*θ*) is the cost, equal to the sum of the normalized residual over all measured time points, *t*; *p* is the parameters; *y*_*t*_ is the measured data and *ŷ*_*t*_(*θ*) is the model simulations; *SEM*_*t*_ is the standard error of the mean for the measured data.

### Statistical analysis

To reject models, we used the *χ*^2^-test with a significance level of 0.05. We used 137 degrees of freedom for the original training data (144 data points, minus 7 scaling parameters) leading to a threshold for rejection of *χ*^2^(0.05, 137) ≈ 165.3. For the extended set of experimental data, used after diabetes was introduced (from the section **Estimating the extent of altered reesterification in type 2 diabetes** in the results), we used 152 degrees of freedom, resulting in a threshold for rejection of *χ*^2^(0.05, 152) ≈ 181.8. Any combination of model and parameter set that results in a cost (Eq. (12)) above the threshold must be rejected. If no parameter set exists for a model that results in a sufficiently low cost, the model structure must be rejected.

### Uncertainty estimation

The uncertainty of both the parameters and the model simulations for estimation, validation, and predictions, were gathered as proposed in [23] and implemented in [14]. In short, the desired property (either a parameter value or a single simulated value) was either maximized or minimized, while requiring the cost (Eq. (12)) to be below the *χ*^2^-threshold. In traditional profile-likelihood analysis (Eq. (13)),

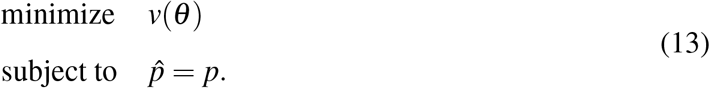

the cost *v*(*θ*) is minimized while the property 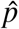is fixed to a value *p* and stepped through to find the boundaries of the property. Here, we instead inverse the problem and solve it directly, see Eq. (14):

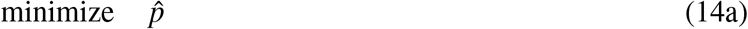

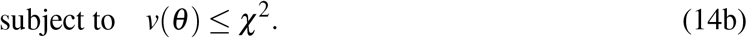

In practice, the constraint (Eq. (14b)) is relaxed into the objective function as a L1 penalty term with an offset if *V* (*θ*) > *χ*^2^.

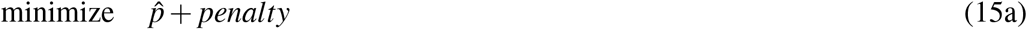

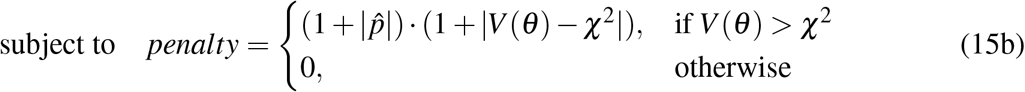

We solved the maximization problem as a minimization problem (same as in Eq. (15)), by changing sign of the property in the objective function to 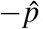.

### Optimization and software

We used MATLAB R2020a (MathWorks, Natick, MA) and the IQM toolbox (IntiQuan GmbH, Basel, Switzerland), a continuation of [24], for modelling. The parameter values were estimated using the enhanced scatter search (eSS) algorithm from the MEIGO toolbox [25]. eSS were restarted multiple times, partially run in parallel at the local node of the Swedish national supercomputing centre (NSC). We allowed the parameter estimation to freely find the best possible combinations of parameter values, within boundaries. The bounds of the parameter values are given in supplementary Table S2.

## Data and model availability

The experimental data as well as the complete code for data analysis and modelling are available at https://github.com/willov/lipolysis (DOI: 10.5281/zenodo.5070401) and is mirrored at https://gitlab.liu.se/ISBgroup/projects/lipolysis.

## Author Contributions

All authors were involved in the writing of the paper. EN designed the study. WL, JE, and EN performed the modelling. All authors analysed data.

## Abbreviations

cAMP: cyclic AMP
IR: insulin receptor
PKB: protein kinase B
PDE3B: phosphodiesterase 3B
AR: adrenergic receptor
HSL: hormone-sensitive lipase
AC: adenylyl cyclase
ODE: ordinary differential equation

## Supplemental Material

**Figure S1:**
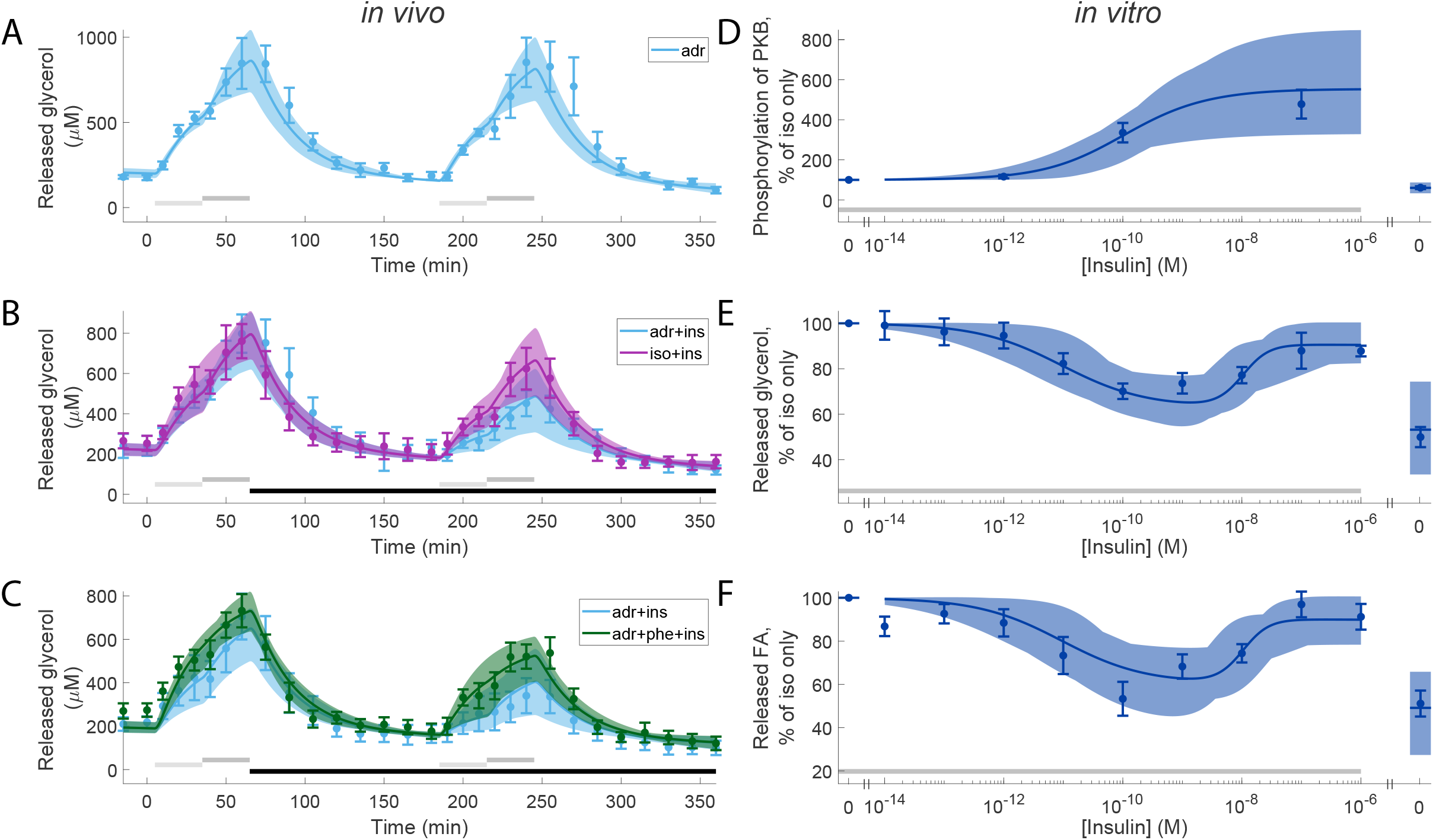
Model agreement with experimental data, with overlaid *in vivo* experiments. In all panels, solid lines represent the model simulation with the best agreement to data, the shaded areas represent the model uncertainty, and experimental data point are represented as mean values with error bars (SEM). (A-C), *in vivo* time-series experiments. (D-F), *in vitro* dose-response experiments. In all subfigures, horizontal bars indicate where stimulations were given. In (A-C), light/dark grey bars indicate low/high adrenergic stimulus (1/10 µM adrenaline or 0.1/1 µM isoproterenol) with or without phentolamine (phe; 100 µM), black bars indicate stimulation with insulin (0.6 nM). In (D-F) grey bars indicate stimulation with isoproterenol (10 nM). In the *in vivo* experiments, experiments with adrenaline are shown in light blue (A-C), with isoproterenol in purple (B), and with the combined stimulation with adrenaline and phentolamine in green (C). In the *in vitro* experiments (D-F), increasing doses of insulin were given together with 10nM isoproterenol in all points except one. The point without isoproterenol got no stimulus and is shown to the right in the graphs.

**Table S1:**
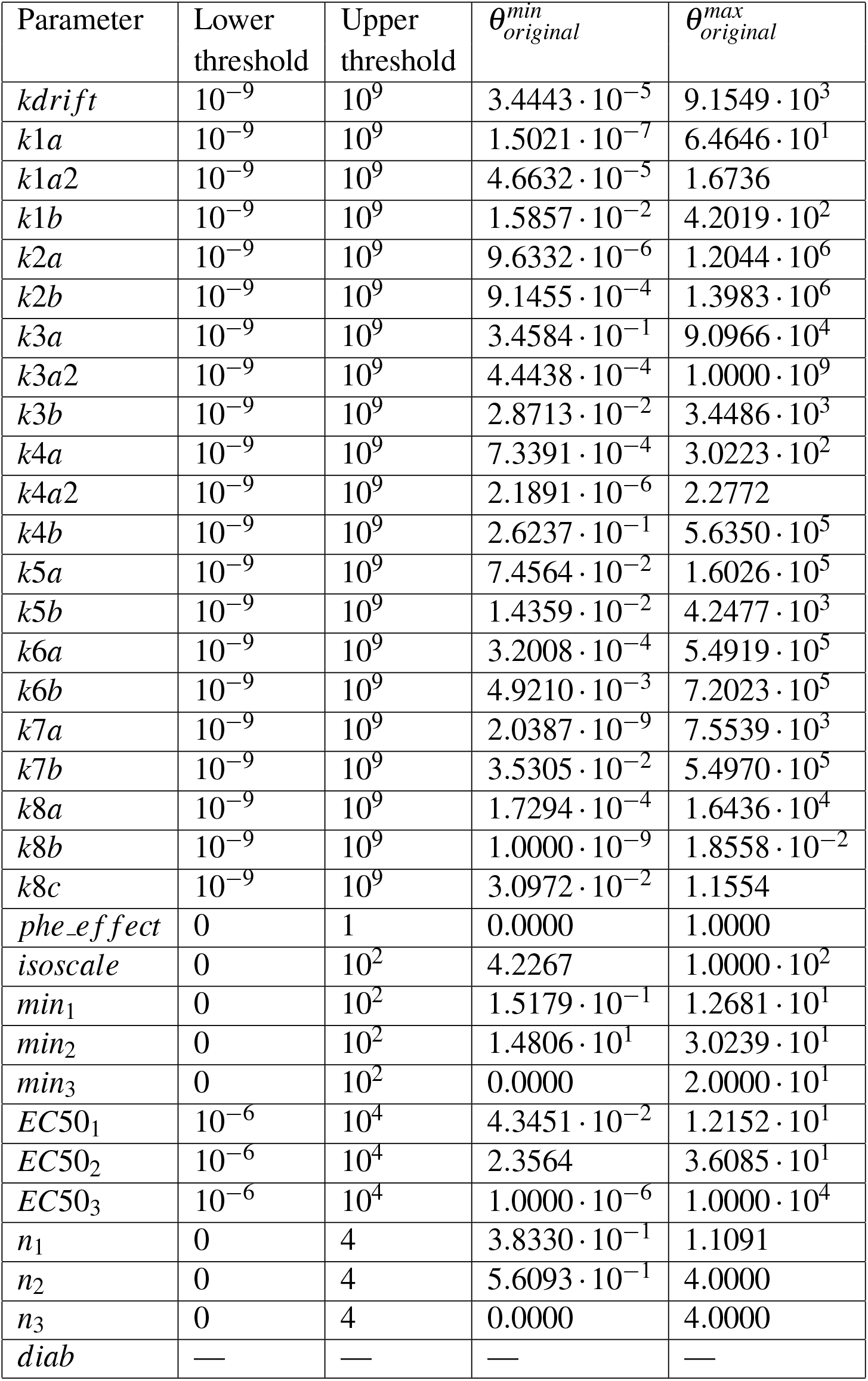
All bounds and estimated values for the free parameters. The parameters were allowed to vary in the range given in Table S2. For the specific parameter being investigated, the bound was relaxed and the threshold for when a parameter was deemed nonidentifiable was set to the value given in the table in columns Lower threshold and Upper threshold). The minimum and maximal found values of a parameter is given in columns 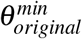, and 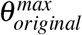 respectively.

**Table S2:** Bounds used for optimization of the free parameters, and the sets of optimal values. The rate parameters (*kx*) were given a free range (10^−6^ to 10^6^). *isoscale* was allowed a 20% deviation from the expected value of 10. For the input functions, the minimum values *min*_*x*_ was given a range from zero to 20% of max, the steepness *n*_*x*_ was given a range from 0 to 2, and the *EC*50_*x*_ was given a free range for all doses used in the dataset from [3] (10^−5^ to 10^3^ nM), except for *EC*50_1_ which was limited based on the EC50 of IR in [5]. 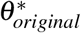 corresponds to the optimal parameter set for the original dataset. 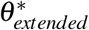 corresponds to the optimal parameter set for the extended dataset.

| Parameter | Lower bound | Upper bound | $\theta^*_{original}$ | $\theta^*_{extended}$ |
| --- | --- | --- | --- | --- |
| *kdrift* | $10^{-6}$ | $10^6$ | 1.8649 | 2.6525 |
| *k1a* | $10^{-6}$ | $10^6$ | $2.5929 \cdot 10^{-2}$ | $8.0752 \cdot 10^{-2}$ |
| *k1a2* | $10^{-6}$ | $10^6$ | $2.7345 \cdot 10^{-2}$ | $1.2840 \cdot 10^{-2}$ |
| *k1b* | $10^{-6}$ | $10^6$ | $6.1169 \cdot 10^{-1}$ | $2.0825 \cdot 10^{-1}$ |
| *k2a* | $10^{-6}$ | $10^6$ | 6.5912 | $1.2136 \cdot 10^{-2}$ |
| *k2b* | $10^{-6}$ | $10^6$ | $2.0591 \cdot 10^1$ | $9.4937 \cdot 10^{-3}$ |
| *k3a* | $10^{-6}$ | $10^6$ | $9.2000 \cdot 10^1$ | $1.2626 \cdot 10^2$ |
| *k3a2* | $10^{-6}$ | $10^6$ | 9.4628 | $3.0140 \cdot 10^1$ |
| *k3b* | $10^{-6}$ | $10^6$ | $1.1379 \cdot 10^1$ | $1.4944 \cdot 10^1$ |
| *k4a* | $10^{-6}$ | $10^6$ | $8.1360 \cdot 10^{-1}$ | 1.1020 |
| *k4a2* | $10^{-6}$ | $10^6$ | $3.9464 \cdot 10^{-3}$ | $1.5750 \cdot 10^{-2}$ |
| *k4b* | $10^{-6}$ | $10^6$ | $5.6524 \cdot 10^2$ | $1.2609 \cdot 10^2$ |
| *k5a* | $10^{-6}$ | $10^6$ | $2.6695 \cdot 10^2$ | $2.8775 \cdot 10^1$ |
| *k5b* | $10^{-6}$ | $10^6$ | 3.0725 | 2.0128 |
| *k6a* | $10^{-6}$ | $10^6$ | $4.0644 \cdot 10^{-2}$ | $4.1938 \cdot 10^{-1}$ |
| *k6b* | $10^{-6}$ | $10^6$ | $8.9551 \cdot 10^{-3}$ | $2.4450 \cdot 10^{-1}$ |
| *k7a* | $10^{-6}$ | $10^6$ | $2.1389 \cdot 10^{-1}$ | $9.7287 \cdot 10^{-2}$ |
| *k7b* | $10^{-6}$ | $10^6$ | $5.6389 \cdot 10^1$ | $1.8381 \cdot 10^2$ |
| *k8a* | $10^{-6}$ | $10^6$ | $6.5756 \cdot 10^1$ | $2.6920 \cdot 10^3$ |
| *k8b* | $10^{-6}$ | $10^6$ | $5.4205 \cdot 10^{-3}$ | $2.1333 \cdot 10^{-2}$ |
| *k8c* | $10^{-6}$ | $10^6$ | $3.8648 \cdot 10^{-2}$ | $3.4005 \cdot 10^{-2}$ |
| *phe_effect* | 0.6 | 1 | $8.8329 \cdot 10^{-1}$ | $8.4505 \cdot 10^{-1}$ |
| *isoscale* | 8 | $1.2 \cdot 10^1$ | 9.4266 | 8.0000 |
| *min<sub>1</sub>* | 0 | $2 \cdot 10^1$ | 1.9994 | 1.1158 |
| *min<sub>2</sub>* | 0 | $2 \cdot 10^1$ | $2.00000 \cdot 10^1$ | $2.0000 \cdot 10^1$ |
| *min<sub>3</sub>* | 0 | $2 \cdot 10^1$ | 0.0000 | 0.0000 |
| *EC50<sub>1</sub>* (nM) | 0.5 | 1.1 | 1.2454 | 1.2454 |
| *EC50<sub>2</sub>* (nM) | $10^{-5}$ | $10^3$ | $1.2853 \cdot 10^1$ | $1.3356 \cdot 10^1$ |
| *EC50<sub>3</sub>* (nM) | $10^{-5}$ | $10^3$ | 2.8643 | 4.1559 |
| *n<sub>1</sub>* | 0.5 | 2 | $6.1281 \cdot 10^{-1}$ | $6.6829 \cdot 10^{-1}$ |
| *n<sub>2</sub>* | 0.5 | 2 | 1.9807 | 1.7424 |
| *n<sub>3</sub>* | 0.5 | 2 | 2.0000 | 2.0000 |
| *diab* | 0 | 1 | — | $9.4622 \cdot 10^{-1}$ |

**Figure S2:**
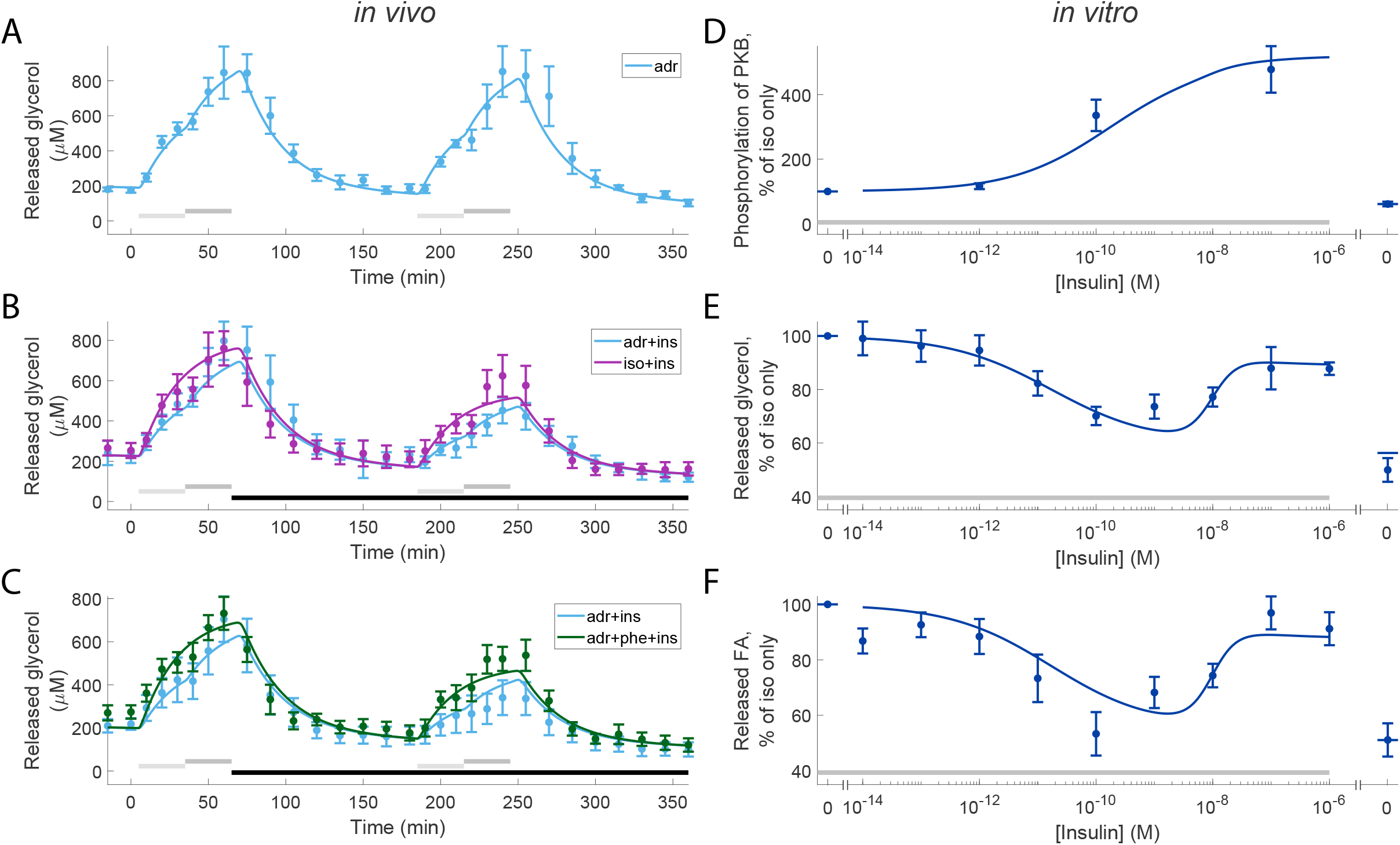
Model agreement with experimental data, with overlaid *in vivo* experiments, for the model without insulin action-3. In all panels, solid lines represent the model simulation with the best agreement to data, the shaded areas represent the model uncertainty, and experimental data point are represented as mean values with error bars (SEM). (A-C), *in vivo* time-series experiments. (D-F), the *in vitro* dose-response experiments. In all subfigures, horizontal bars indicate where stimulations were given. In (A-C), light/dark grey bars indicate low/high adrenergic stimulus (1/10 µM adrenaline or 0.1/1 µM isoproterenol) with or without phentolamine (phe; 100 µM), black bars indicate stimulation with insulin (0.6 nM). In (D-F) grey bars indicate stimulation with isoproterenol (10 nM). In the *in vivo* experiments, experiments with adrenaline are shown in light blue (A-C), with isoproterenol in purple (B), and with the combined stimulation with adrenaline and phentolamine in green (C). In the *in vitro* experiments (D-F), increasing doses of insulin were given together with 10nM isoproterenol in all points except one. The point without isoproterenol got no stimulus and is shown to the right in the graphs.

**Figure S3:**
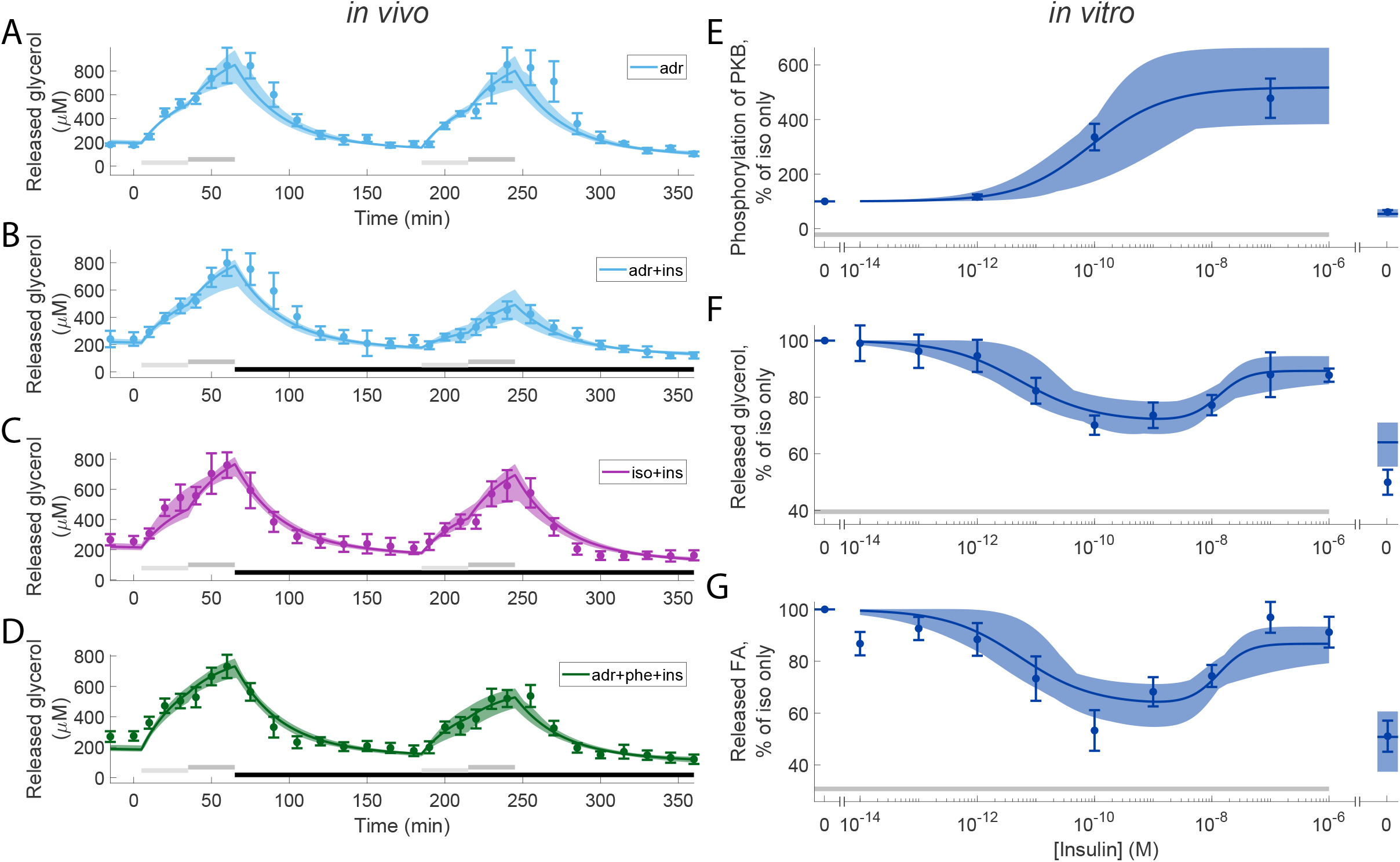
Model agreement with the extended set of experimental data not shown in Fig. 4. Estimation data and model simulations, for the original data (e.g. used in Fig. 3). In all panels, solid lines represent the model simulation with the best agreement to data, and experimental data point are represented as mean values with error bars (SEM). (A-D), *in vivo* time-series experiments. (E-G), *in vitro* dose-response experiments. In all subfigures, horizontal bars indicate where stimulations were given. In detail, light/dark grey bars indicate stimulation with: 1/10 µM adrenaline in (A,B), 0.1/1 µM isoproterenol in (C), and 1/10 µM adrenaline with 100 µM phentolamine. Black bars in (B-D) indicates stimulation with 0.6 nM insulin. In (E-G) grey bars indicate stimulation with isoproterenol (10 nM). In the *in vivo* experiments, experiments with adrenaline are shown in light blue (A-C), with isoproterenol in purple (B), and with the combined stimulation with adrenaline and phentolamine in green (C). In the *in vitro* experiments (D-F), increasing doses of insulin were given together with 10 nM isoproterenol in all points except one. The point without isoproterenol got no stimulus and is shown to the right in the graphs.

**Figure S4:**
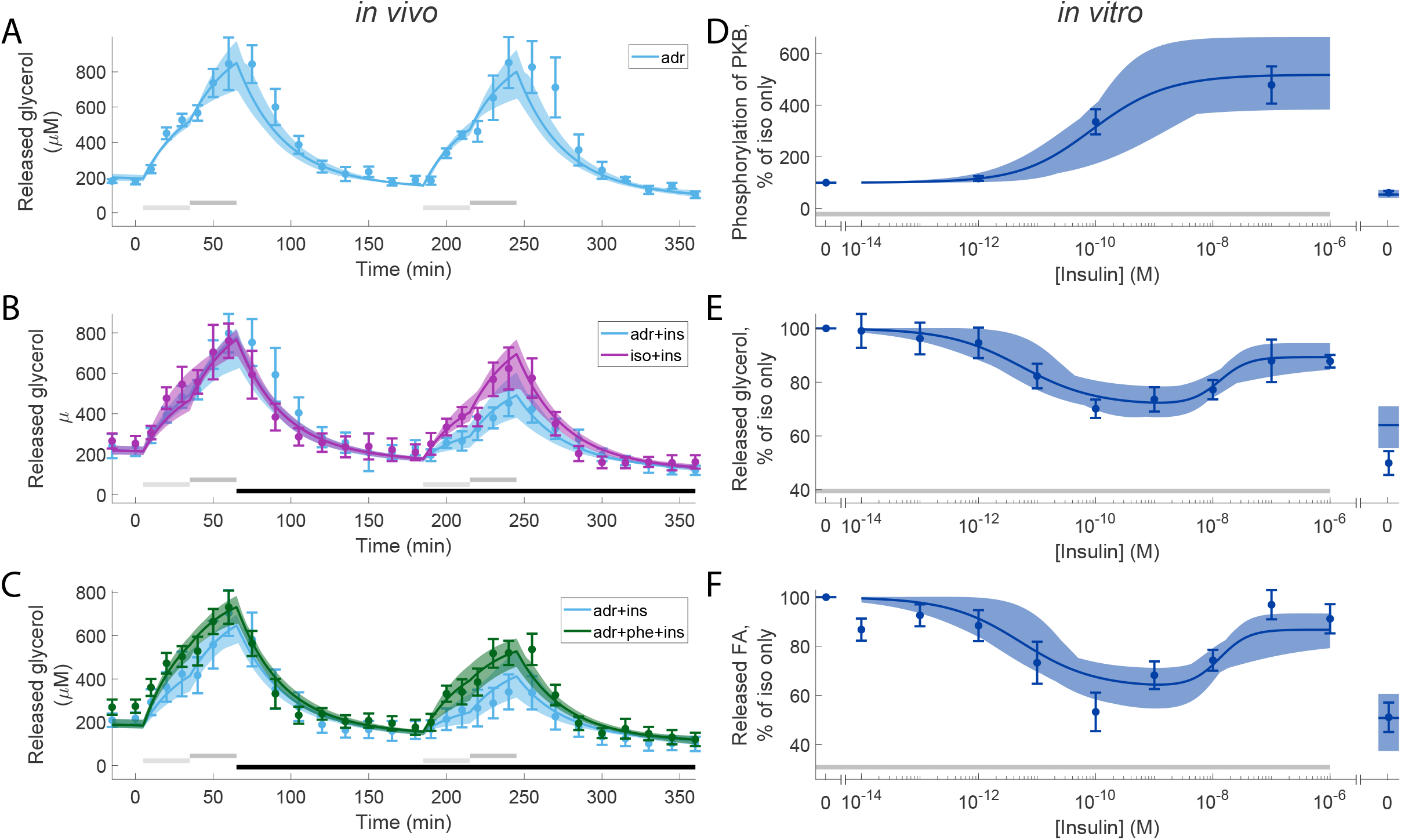
Model agreement with the extended set of experimental data not shown in Fig. 4, with overlaid *in vivo* experiments. Estimation data and model simulations, for the original data (e.g. used in Fig. 3). In all panels, solid lines represent the model simulation with the best agreement to data, the shaded areas represent the model uncertainty, and experimental data point are represented as mean values with error bars (SEM). (A-C), *in vivo* time-series experiments. (D-F), *in vitro* dose-response experiments. In all subfigures, horizontal bars indicate where stimulations were given. In (A-C), light/dark grey bars indicate low/high adrenergic stimulus (1/10 µM adrenaline or 0.1/1 µM isoproterenol) with or without phentolamine (phe; 100 µM), black bars indicate stimulation with insulin (0.6 nM). In (D-F) grey bars indicate stimulation with isoproterenol (10 nM). In the *in vivo* experiments, experiments with adrenaline are shown in light blue (A-C), with isoproterenol in purple (B), and with the combined stimulation with adrenaline and phentolamine in green (C). In the *in vitro* experiments (D-F), increasing doses of insulin were given together with 10nM isoproterenol in all points except one. The point without isoproterenol got no stimulus and is shown to the right in the graphs.

